# Integrated decoding hematopoiesis and leukemogenesis at single-cell resolution and its clinical implication

**DOI:** 10.1101/2020.02.21.960401

**Authors:** Pengfei Qin, Yakun Pang, Wenhong Hou, Ruiqing Fu, Yingchi Zhang, Xuefei Wang, Guofeng Meng, Qifa Liu, Xiaofan Zhu, Ni Hong, Tao Cheng, Wenfei Jin

**Author notes:** These authors contributed equally to this work (Pengfei Qin, Yakun Pang, Wenhong Hou, Ruiqing Fu, Yingchi Zhang, Xuefei Wang).

## Abstract

Integrated analyses of hematopoiesis and leukemogenesis could provide new biological and clinical insights^1,2^. Here we constructed a comprehensive cell atlas using ∼100,000 single cell transcriptomes of bone marrow mononuclear cells and its subsets from 5 healthy samples and 13 leukemia samples. Trajectory analysis showed hematopoietic stem cells continuously differentiated through tree-like structure into 7 lineages. Although our single cell analyses showed the major cellular heterogeneity were inter-patients, there were distinct leukemia cell subpopulations within patient. We developed counterpart composite index (CCI) for projecting leukemia cell subpopulation into reference hematopoietic cells, to identify their healthy counterparts that could predict leukemia subtype and clinical outcome. Interestingly, we found the trajectories of leukemogenesis were similar to that of their healthy counterparts, indicating lineage relationship of leukemia cell subpopulations. Single cell analyses of leukemia patient at diagnosis, refractory, remission and relapse vividly presented dynamics of cell population and the underlying genes. Single cell transcriptomic variants analyses showed the relapsed leukemia cells, whose healthy counterparts were closer to the root of hematopoietic tree, were derived from an early minor leukemia cell population. This study not only increased our understanding of hematopoiesis and leukemogenesis, but also provided an approach for leukemia classification and clinical outcome prediction.

## Main

Single cell RNA sequencing (scRNA-seq) provides unbiased gene expression profile of individual cells that is highly complementary to the immunological phenotypes approaches^3^. Recent massively parallel scRNA-seq enabled routine analyses of thousands of single cells for inferring the developmental trajectories^4–8^. In particular, single cell analysis of hematopoietic stem and progenitor cells (HSPCs) demonstrated hematopoiesis is a continuous process rather than discrete stepwise process ^8–12^. However, inconsistency persists among those studies, e.g. Velten et al. proposed the CLOUD-HSPCs model in which HSPCs directly gave rise to distinct lineage-committed populations ^9^, while Tusi et al. proposed a continuously hierarchies model ^8^. Furthermore, the molecular mechanisms underlying hematopoietic lineage commitment remain poorly understood ^10^.

Leukemia patients are still under the threat of relapse and drug resistance due to leukemia heterogeneity. Analyses of cancer stem cell in chronic myeloid leukemia identified distinct subpopulations of therapy-resistant stem cells ^13^. Recent single cell study on acute myeloid leukemia (AML) revealed primitive leukemia cells aberrantly co-expressed stemness and myeloid priming genes ^1^. However, our knowledge about the relationship of leukemia cell subpopulations and dynamic progression of leukemia are still limited. In this study, we constructed a comprehensive cell atlas of hematopoietic cells, and a hierarchically continuous transition model for hematopoiesis. We developed CCI to project leukemia cells to reference hematopoietic cells and identified the healthy counterparts of leukemia cell, which could predict leukemia subtype and clinical outcome. Single cell RNA-seq analysis of a patient at diagnosis, refractory, remission and relapse vividly demonstrated dynamics of leukemia progression.

## Results

### A comprehensive cell atlas of healthy BMMCs

Bone marrow is the primary tissue for blood cell production, and generates hundreds of billions of blood cells and immune cells per day. In order to gain further biological insights into hematopoiesis and leukemogenesis, we established approaches and pipelines for single cell analysis of BMMCs and its subsets from 5 healthy samples and 13 leukemia samples (**Fig. 1a, Extended Data Fig. 1a-1c and Table S1**). The 18,751 cells from 4 healthy BMMCs were clustered into 17 distinct cell clusters using Louvain clustering algorithm and visualized by t-Distributed Stochastic Neighbor Embedding (tSNE) ^14^ (**Fig. 1b**). We identified the cell type of each cluster based on their specific highly expressed genes (**Fig. 1c, 1d**). The frequencies of major identified cell types in BMMCs essentially consists with the expectations (**Fig. 1b-1d**): 1.39% HSPCs (AVP+ and CD34+), 10.15% erythroid progenitor cells (EPCs) (GYPA+ and KLF1+), 0.08% megakaryocytes (Mk) (PF4+ and GP9+), 1.25% myelocytes (ELANE+ and MPO+), 2.91% monocytes (LYZ+ and CD14+), 17.67% B cells (CD79A+), 53.11% T cells (CD3D+), 11.62% natural killer cells (NK) (FCGR3A+ and NCAM1+), 0.04% stromal cells (CXCL12+ and COL6A1+). Furthermore, the frequencies of cell types were highly consistent across different samples (r^2^=0.96; **Extended Data Fig. 1e**).

**Fig. 1.**
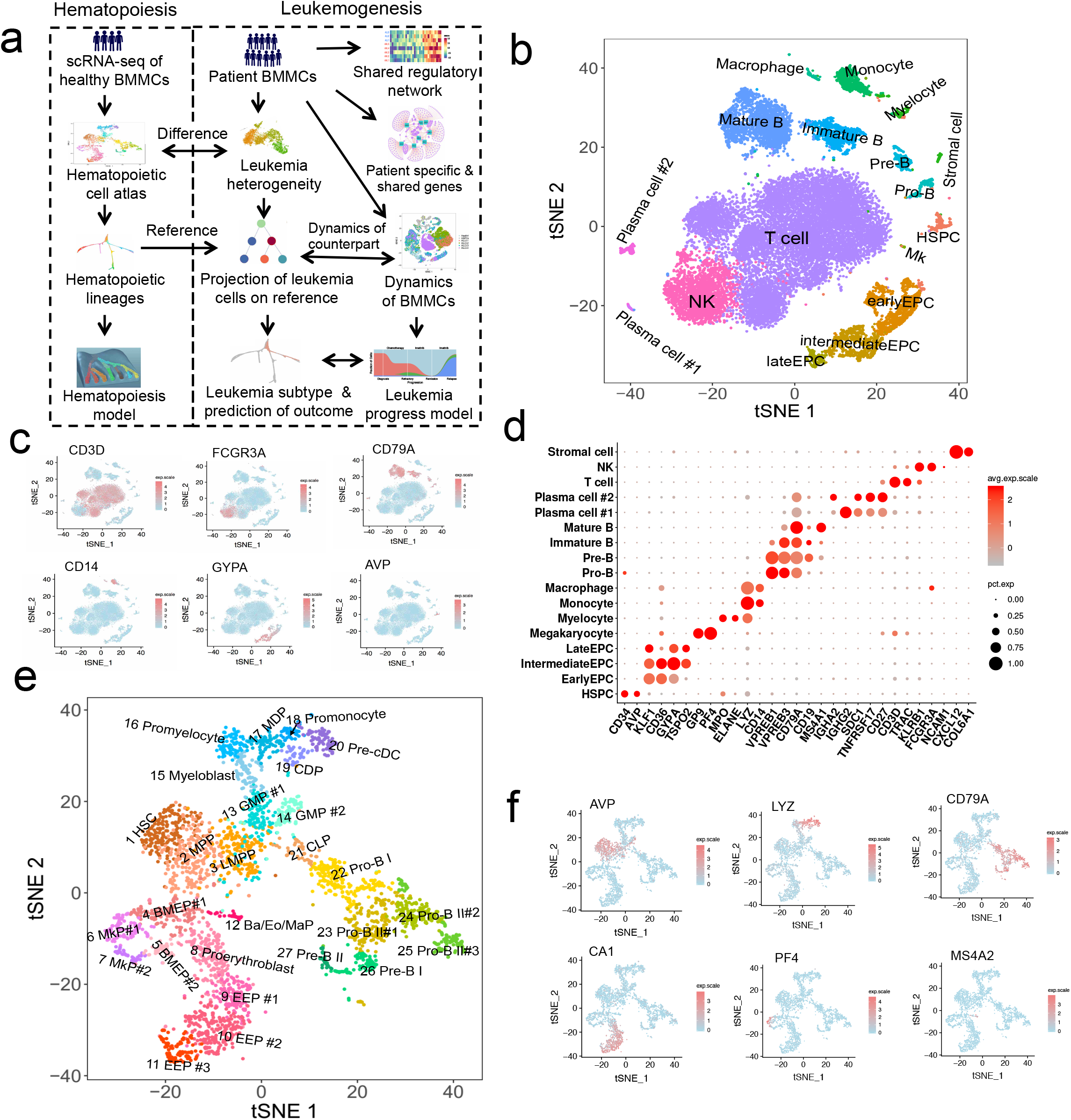
The cell atlas of bone marrow mononuclear cells (BMMCs) and HSPCs (CD34+) cells in healthy individuals. **a.** The schematic of this study. **b.** tSNE projection of BMMCs, colored by inferred cell type. **c.** tSNE projection of BMMCs with each cell colored based on their normalized expression of CD3D, FCGR3A, CD79A, CD14, GYPA and AVP, respectively. **d.** Normalized expression level and expression percentage of the cell type specific genes in 17 cell populations in BMMCs. **e.** tSNE projection of HSPCs (CD34+) cells, colored by inferred cell type. **f.** tSNE projection of HSPCs with each cell colored based on their normalized expression of AVP, LYZ, CD79A, CA1, PF4 and MS4A2.

Interestingly, we identified two distinct plasma cell populations both highly expressed SDC1, CD27 and TNFRSF17 (**Fig. 1b, 1d**). Plasma cell #1 highly expressed IGHG1, IGHG2, IGHG3, IGHG4 and IGKC, while plasma cell #2 highly expressed IGHA1, IGHA2 and translation associated genes (**Fig. 1d, Extended Data Fig. 1d**), indicating their different cellular status and functions. We further identified a lot T cell subset in BMMCs (**Extended Data Fig. 1f**). Compared with peripheral blood mononuclear cells (PBMCs) ^15^, BMMCs contain much more subpopulations and are more active (**Extended Data Fig. 1g-1j**).

### HSPCs form a single connected entity on tSNE projection

Due to the limited number of HSPCs in BMMCs (**Fig. 1b**), CD34+ cells, representing HSPCs, were enriched by fluorescence-activated cell sorting (FACS) for investigating hematopoiesis **(Extended Data Fig. 2a**). The HSPCs essentially formed a single connected entity extending in several directions on tSNE projection and were clustered into 27 clusters for better understanding of the hematopoietic process (**Fig. 1e**). We inferred the cell type of each cluster by checking the expression of hematopoietic lineage specific genes (**Fig. 1e, Extended Data Fig. 2b**), such as HSC (EMCN+, THY1+, MEG3+, HES1+, Lin-; cluster 1), megakaryocytic progenitors (MkP) (PF4+, GP9+; clusters 5, 7), early erythroid progenitors (EEP) (APOE+, CD36+, CA1+; clusters 8-11), neutrophil, monocyte and DC progenitors (CSF3R+, MPO+ and LYZ+; clusters 13-20) (**Fig. 1f, Extended Data Fig. 2b**), lymphoid progenitors (CD79A+, IGHM+, VPREB1+; clusters 21-27) (**Fig. 1F and Extended Data Fig. 2b**)

Cluster 12 highly expressed mast cell and basophil specific genes including HDC, TPSAB1 and MS4A2, as well as eosinophil specific genes including PRG2 and CPA3 (**Fig. 1f and Extended Data Fig. 2b**), which could be recently reported Basophil/Eosinophil/Mast progenitors (Ba/Eo/MaP) ^10,16^. We did not detect any cluster with gene expression patterns similar to common myeloid progenitor (CMP) (CD34+, CD38+, CD123+, CD45RA-, CD10- and Lin-), consistent with recent studies showing that CMP is a heterogeneous mixture of erythroid and myeloid primed progenitors ^10,17,18^. Moreover, we observed multiple subpopulations within pre-defined MkP, EEP, GMP, Pro-B II and so on (**Fig. 1e**). The expression levels of many genes are gradually changing along the three EEP populations, among which the expression levels of HBA1, TFRC, GYPA, ALAS2, PLK1 and MKI67 gradually increased as the distance to HSC increase (**Extended Data Fig. 2b, 2i, 2j**). Overall, HSPCs contain a substantial higher fraction of cells in active cell cycles and cell states than that of other BMMCs (**Extended Data Fig. 2c-2h**). Interestingly, major early stem and progenitor cells (HSC, MPP, LMPP) are in resting phase while major later progenitors are in active proliferation (**Extended Data Fig. 2e, 2f**), potentially indicating the early progenitor constitutes the major cell pool for regulating hematopoiesis while later progenitors are in simple transitional states.

### Continuous hematopoietic lineages with hierarchical structure

We implemented Slingshot ^19^ and SPRING ^20^ on HSPCs to conduct pseudo-time inference. Pseudo-time ordering of HSPCs exhibits a tree-like structure in which HSC forms the root, from which seven lineages gradually emerged with a hierarchical structure (**Fig. 2a, 2b, 2c**), essentially consistent with the cell lineages base on PCA projection (**Extended Data Fig. 3a**). The results are consistent with recent reports that hematopoiesis is a continuous process ^8–10^, but with different structure and process. In particular, HSC was embedded between Mk lineage and Lym lineage in Tusi *et al.* ^8^, potentially caused by limited number of early HSPCs in their study. Clusters 4-5 derived from HSC/MPP and are progenitors of Ba/Eo/Ma lineage, Mk lineage and erythroid (Ery) lineage (called BMEP), which are quite different from classic hematopoietic structure. This study also showed neutrophil lineage was derived from GMP while Ba/Eo/Ma lineage was derived from BMEP, different from classic hematopoietic model that granulocytes shared common progenitor ^21,22^.

**Fig. 2.**
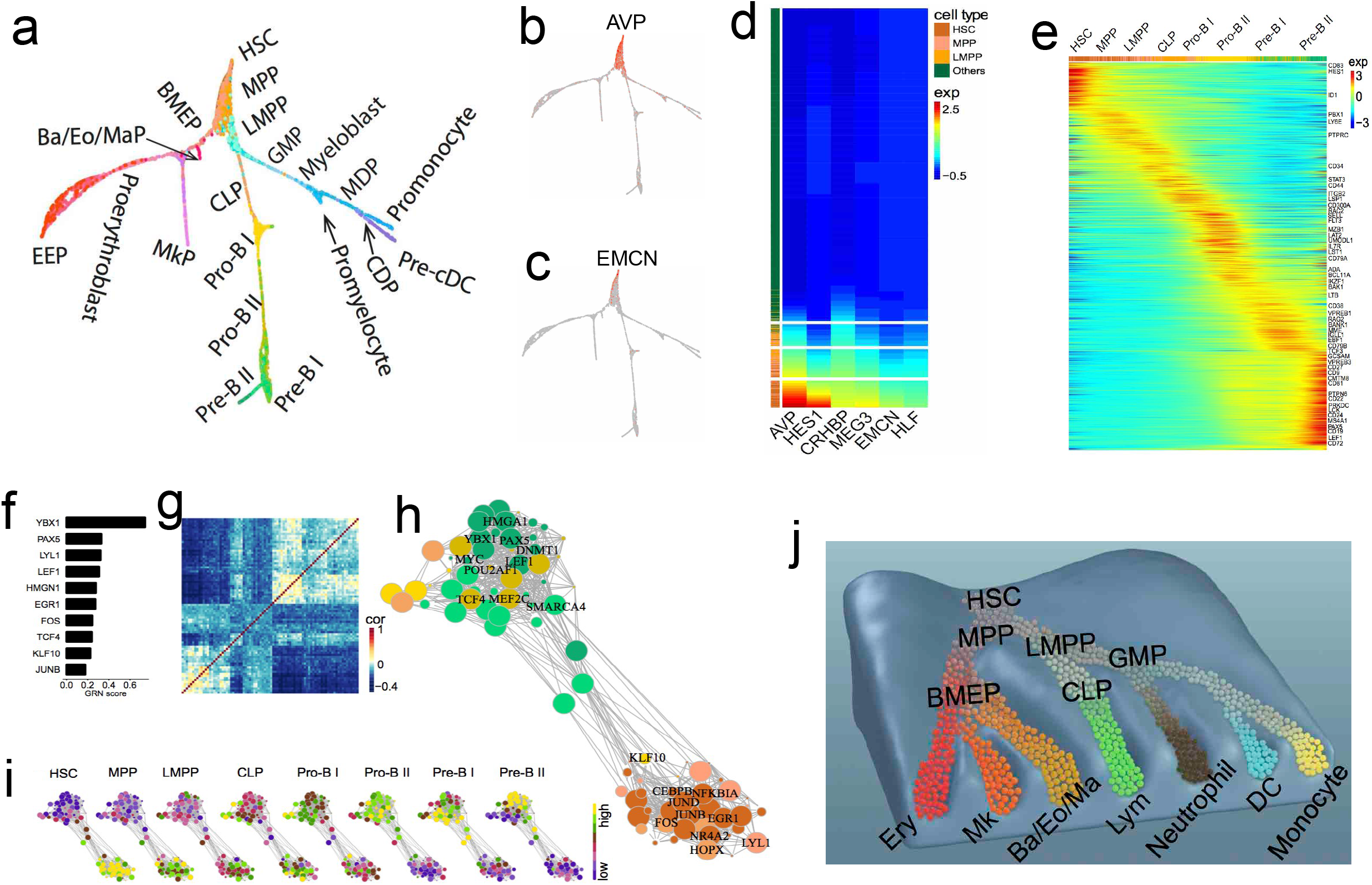
Hematopoietic cell lineages and hierarchically continuous transition model for hematopoiesis. **a**. Hematopoietic lineages visualized by SPRING, cells were colored by cell type as in Fig. 1e. **b-c**. Expressions of AVP (b) and EMCN (c) are decreasing along hematopoietic lineages. **d.** Heatmap of normalized expression level of early hematopoietic markers along lineages. **e.** Heatmap of transcriptomic dynamics during lymphopoiesis. **f-i.** Coordinated TFs and networks underlying lymphoid lineage. The top 10 coordinated TFs (f); correlation of the coordinated TFs g); network of coordinated TFs, in which the size of each node represents magnitude of expression (h); dynamics of TFs expression in regulatory networks along lymphoid lineage (i), in which each node was colored by average expression. **j**. Hierarchically continuous transition model for hematopoiesis.

Heatmap analysis showed the expression of early hematopoietic marker genes, such as AVP, HES1, CRHBP, MEG3, EMCN and HLF, are gradually decreasing along pseudo-time (**Fig. 2d**). Although cells were clustered into different populations, their expressions do not show significant changes across the boundaries, further supporting gradually decrease of stemness along hematopoietic lineage. Attenuations of expression vary greatly from gene to gene, and different cells showed different gene expression patterns (**Fig. 2d**), indicating each cell with a unique status during hematopoiesis. Furthermore, by integrating HSPCs and BMMCs together (**Extended Data Fig. 3b**), we constructed a much comprehensive cell atlas and hematopoietic lineages (**Extended Data Fig. 3c, 3d**), in which HSPC located in the center while erythrocytes, lymphocytes and monocytes from BMMCs extended at terminal branch (**Extended Data Fig. 3e**).

### Lineage-coordinated genes, transcription factors (TFs) and networks

We identified thousands of lineage-coordinated genes with expression gradually shifting along hematopoietic lineages (**Fig. 2e; Extended Data Fig. 4a, 4b**). For instance, the expression level of CD79A, VPREB3 and PAX5 are increasing along the lymphoid (Lym) lineage (Fig. 2e). The observation that many genes changed continuously along the lineages further supports our presumption of hematopoiesis being a continuous process. We further identified lineage coordinated TFs of each lineage using gene regulatory networks (GRN) scores ^23^. Among the top 10 Lym lineage coordinated TFs (**Fig. 2f**), PAX5, LYL1, LEF1, HMGN1, FOS and JUNB have been reported to play important roles in lymphopoiesis ^24^, while YBX1, EGR1, TCF4 and KLF10 are newly identified. We further identified SPL1, ZEB2, CEBPA and IRF1 and IRF8 in DC lineage; SPL1, CEBPD, JUNB, CEBPA and KLF2 in neutrophil lineage; SPL1, JUNB, CEBPA and FOS in monocyte lineage; KLF1, GATA1, ZEB2 and MYC in Ery lineage; ZBTB16, GATA1, ZEB2 and KLF2 in Ba/Eo/Ma cell lineage; FLI1, GATA1, ZEB2 and GFI1B in Mk lineage (**Extended Data Fig. 4c-4h**).

We constructed lineage coordinated TF network based on co-expression network of lineage coordinated TFs. There are two major sub-networks in Lym lineage coordinated TF network with strong intra-subnetwork interactions (**Fig. 2g, 2h**). The subnetwork usage is gradually shifting from the one highly active in HSC to the one highly active in B-cell progenitor along lymphopoiesis (**Fig. 2i**). In contrast, there is only one major connective unit in the neutrophil lineage coordinated TF network, in which TF usage is shifting in the same network during neutrophil genesis (**Extended Data Fig. 4d**). Overall, we observed two or more subnetworks in Ery, Mk, Lym and DC lineages, in which active networks was gradually shifting from one subnetwork to another; while we only observed one major compact core in neutrophil, monocyte and Ba/Eo/Ma lineages with TF usage shifting in the same network (**Extended Data Fig. 4c-4h**), indicating different models for lineage regulations.

### Hierarchically continuous transition model for hematopoiesis

We propose a hierarchically continuous transition model to explain the hematopoiesis process from HSC to distinct hematopoietic lineages (**Fig. 2j**). In this model, the hematopoietic system is a dynamic equilibrium system composed of a large number of transitional states/cells, among which contiguous ones could mutually convert into each other. Thus, the cell states were only affected by its previous states and compensation effect promotes the HSC transition to vacant slots. The cell fate of stem cell is not pre-defined but is gradually determined during the differentiation. Cells sorted by FACS can be treated as a continuum composed of a fraction of transitional states but lost other states. We could consider the classic step-wise model as a specific case of our continuous transition model, in which many transitional states have been missed. The continuous transition model bridges the gap between the classic step-wise models based on FACS sorting and recent continuous models based on single cell sequencing. This model provides an essential reference for understanding of leukemia heterogeneity and leukemogenic process.

### Leukemia heterogeneity and shared features among patients

The molecular heterogeneity of the leukemia has significant impact on leukemia classification and treatment ^25^. To characterize the heterogeneity of leukemia cells, we investigated the single cell transcriptomic data of BMMCs from 8 leukemia patients. tSNE projection showed these cells formed two kinds of clusters, either normal cell clusters with cells from both healthy and patient samples, or patient-specific clusters that only comprise of cells from single patient (**Fig. 3a**). The results indicate leukemia cells were quite different from patient to patient, thus high interpatient heterogeneity. The expressions of hematopoietic lineage specific genes also support the uniqueness of patient-specific leukemia cells (**Fig. 3b**). Compared with healthy BMMCs, the majority of the significantly upregulated gene sets or downregulated gene sets are patient specific, with only a few gene sets being shared among multiple patients (**Fig. 3c, 3d**).

**Fig. 3.**
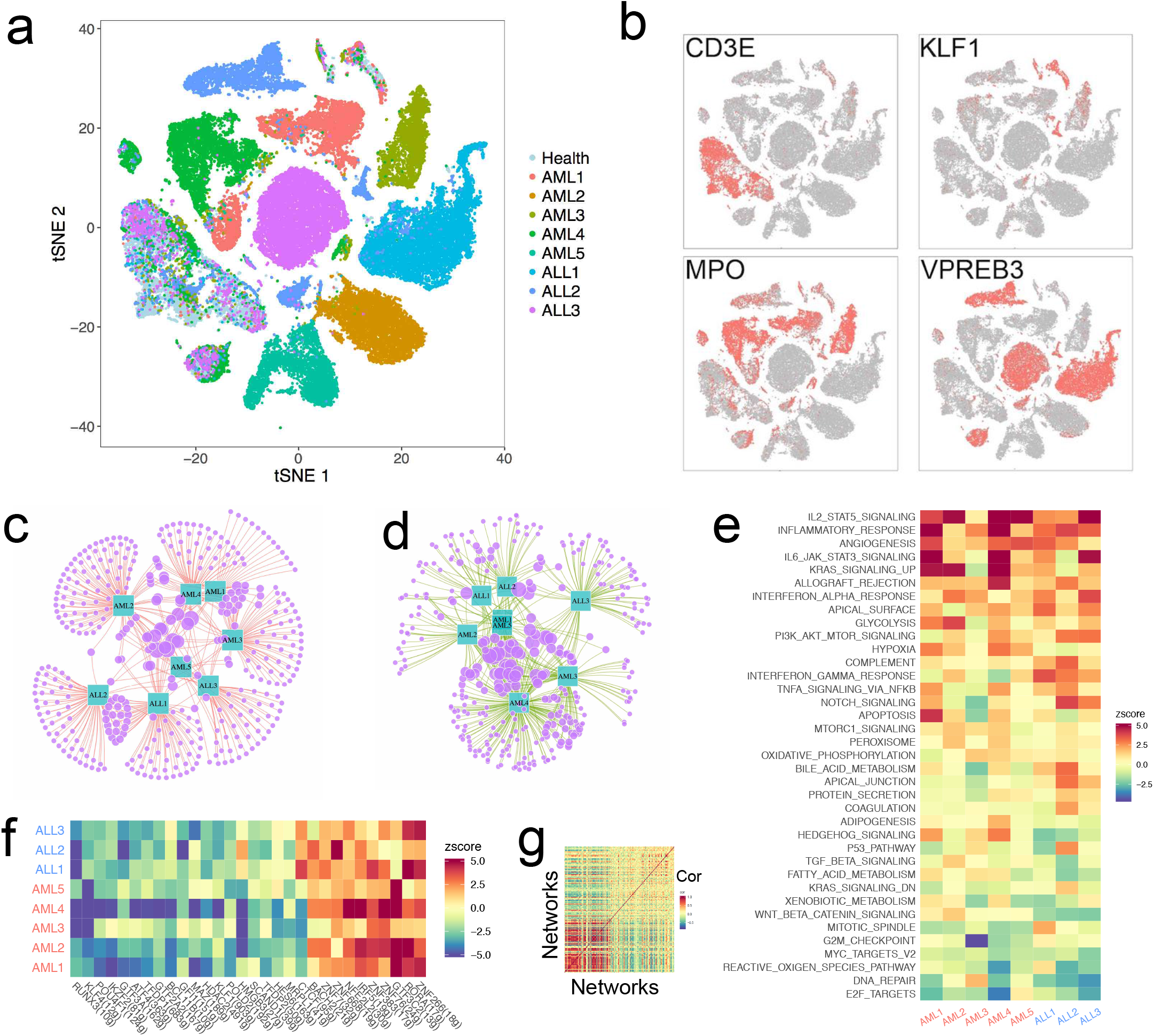
Heterogeneity of leukemia cells and shared features among multiple patients. **a.** t-SNE projection of BMMCs from 1 healthy individual and 8 leukemia patients, colored by different individuals. **b.** t-SNE projection of BMMCs from leukemia patients, with each cell colored based on their expression of CD3E, KLF1, MPO, VPREB3, respectively. **c.** Majority of up-regulated genes among leukemia patients are patient specific. **d.** Majority of down-regulated genes among leukemia patients are patient specific. **e.** Pathway or gene sets commonly up-regulated and down-regulated among leukemia patients. **f.** Shared active and depressed TF networks among leukemia patients. **g.** Correlation of TF networks in leukemia, in which TF networks were sorted based on their activity.

Although leukemia cells exhibited a high heterogeneity among patients, identification of their shared features may provide important clinical implications for diagnosis and treatment. We noticed MIR181A1HG, ITGA4, CD96 and TXNP upregulated in 5 patients and no genes upregulated in all the 8 patients (**Extended Data Fig. 5a**). There are a bunch of gene sets are commonly upregulated or commonly downregulated among all leukemia patients (**Fig. 3e**). The presence of many shared gene sets while absence of shared genes among all patients indicates patients are genetically specific but share signatures/pathways during cancer development. The most significantly upregulated signatures shared by these leukemia patients are IL2-STAT5 signaling, inflammation response, angiogenesis, IL6-JAK-STAT3 signaling, KRAS signaling, allograft rejection and hypoxia, which have been reported in various cancers, indicating these signatures played an important role in cancer development. The most significantly downregulated signatures shared by these leukemia patients are E2F targets, DNA repairs, reactive oxygen species pathway, MYC targets and G2M checkpoint, indicating reduced cell cycle checkpoint and decreased DNA repair activities play an import ant role in leukemia. Indeed, fraction of cells with active cell cycle in leukemia is much higher than that in healthy individual (**Extended Data Fig. 5b-5e**), further indicating higher cell proliferation of leukemia cells.

Compared to healthy BMMCs, we observed the TF networks of ZNF266, RORA, GTF3C2, ZNF76, ZNF383, IRF5 and NFE2L2 being top up-regulated in all leukemia patients (**Fig. 3f**). RORA is the key regulator of embryonic development and cellular differentiation, whose up-regulation may promote the proliferation of the leukemia cells ^26^. IRF5 promotes inflammation by activating genes producing interferons and cytokines ^27,28^. Upregulation of IRF5 network potentially indicates increase of inflammation in leukemia patients, although inflammation alone is inefficient to clean up leukemia cells. The top downregulated TF networks sharing among leukemia patients are RUNX3, KLF4, POU4F1, IKZF2, GTF3A, ATF4, TFDP1, GTF2A2, BCL11B, GFI1, MAZ, HDAC2 and KLF1 (**Fig. 3f**), majority of which are hematopoietic lineage specific, indicating that the healthy hematopoietic process was repressed in these leukemia patients. Heatmap analysis showed that upregulated TF networks and repressed TF networks were correlated, although the correlations of repressed TF networks were weaker than that between upregulated TF networks (**Fig. 3g**).

### Searching healthy counterparts of leukemia cells to predict leukemia subtype and clinical outcome

Heterogeneity of the leukemia cells are associated with disease progression and reaction to chemotherapy ^31,32^. Identifying the leukemia cell subpopulations and inferring their origin could facilitate precise diagnosis and treatments. Here, we developed an approach, called counterpart composite index (CCI), to search for healthy counterpart of each leukemia cell subpopulation by projecting the leukemia cells into reference hematopoietic cell (**Fig. 4a**). In order to improve the statistic power and accuracy, CCI integrates multiple statistics with difference measurements to project leukemia cells to healthy hematopoietic cells (**see METHODs**). Identification of healthy counterparts of leukemia cells not only facilitate our understanding of leukemia progression and leukemia heterogeneity, but also could provide biological insights on the features and functions of the leukemia cell subpopulation via their well-annotated healthy counterparts, thus facilitate prediction of leukemia subtype and clinical outcome.

**Fig. 4.**
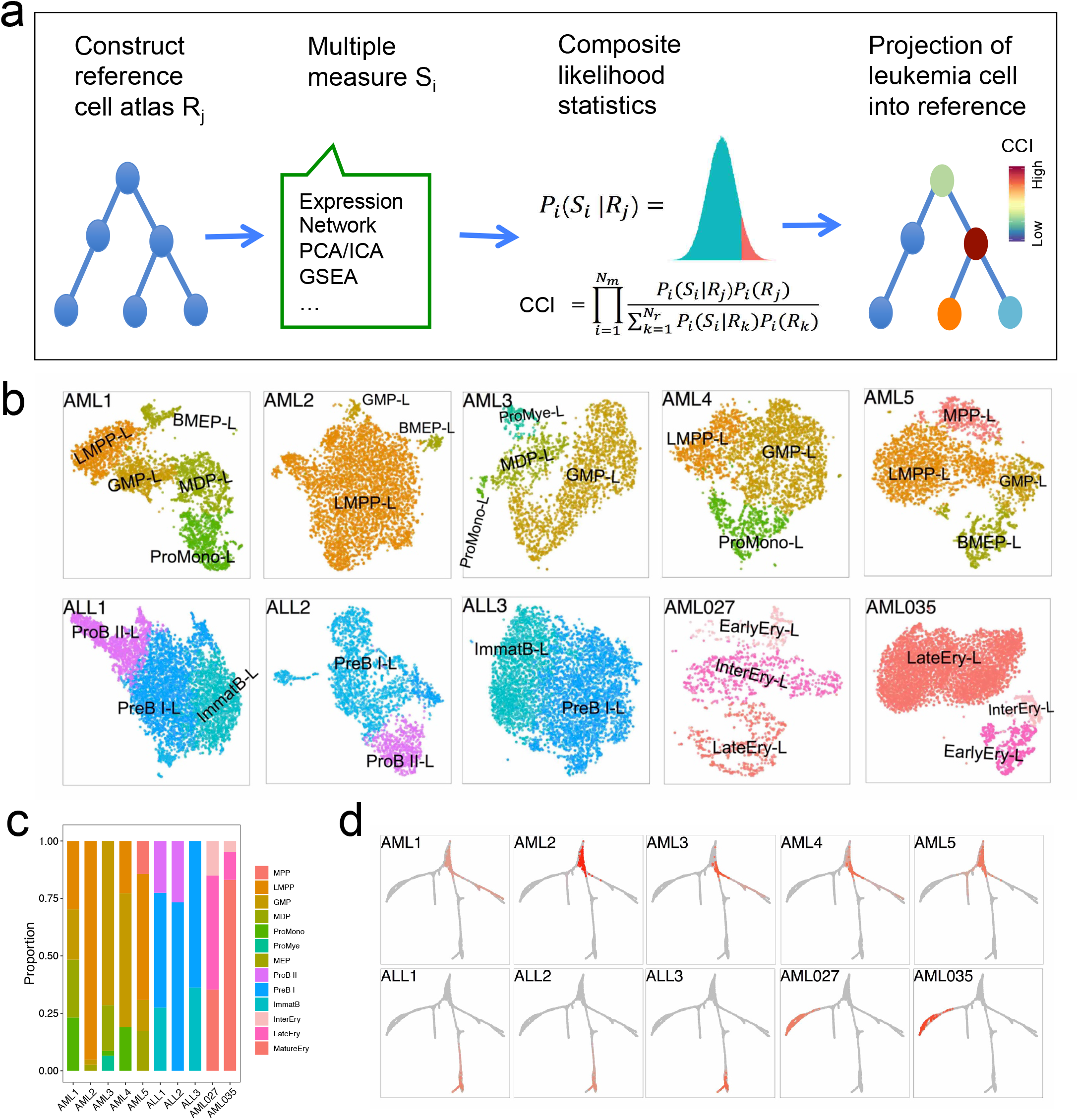
Searching the healthy counterparts of leukemia cells for prediction of leukemia subtype and clinical outcomes. **a.** Schematic of CCI for searching healthy counterparts of leukemia cells. **b.** The healthy counterpart of each leukemia cell subpopulation in the10 leukemia patients, in which the healthy counterparts of leukemia cells from the sample patient are different. Each leukemia cell subpopulation was names after its counterpart by superscript ‘-L’. **c.** Bar plot of leukemia cell subpopulations and their abundance. **d.** Projection of leukemia cells into hematopoietic lineages. Grey indicates reference hematopoietic tree while red indicates leukemia cells. The leukemia cells from ALLs project into lymphoid lineage, while leukemia cells from AMLs project into the non-lymphoid lineages. We found the patient with leukemia cells projecting to the root of the hematopoietic tree had the worst results.

The substructure of leukemia cells is clearly visible and multiple leukemia cell subpopulations could be observed in each patient (**Fig. 4b**). The healthy counterparts of leukemia cell subpopulations were different in the same patient and were different from patient to patient (**Fig. 4b,4c**). The healthy counterparts of leukemia cell in general AML patients include LMPP, BMEP, GMP, MDP, pro-mono, pro-mye and MPP, which belong to HSPCs and myeloid progenitors. The leukemia cell subpopulation was named after its healthy counterpart with ‘-L’ suffix. GMP-L, LMPP-L, BMEP-L are the most common leukemia cell subpopulations in AML patients, among which GMP-L showed up in all the AML patients. The healthy counterparts of leukemia cell from ALL patients include pro-B II, pre-B I, pre-B II and ImmatB, which belong to lymphoid progenitors. PreB I-L showed up in all the three ALL patients and the heterogeneity of leukemia cells in ALL is weaker than that in AML. Interestingly, we found the leukemia cells of two patients (AML027 and AML035) were projected into Ery lineage thus indicated different AML subtypes (**Fig. 4b,4c**). Furthermore, the abundances of leukemia cell subpopulations are quite different from patient to patient (**Fig. 4c**).

Interestingly, projections of the leukemia cells to reference hematopoietic lineages showed the leukemia cells from each patient occupy a flight of hematopoietic lineage (**Fig. 4d**), indicating a similar developmental trajectory between leukemia cells and their healthy counterparts. The leukemia cells from each ALL patient were projected to lymphoid lineage, which are consistent with the clinical diagnosis. The leukemia cells from different AML patients were project into the different myeloid lineages or the root of the hematopoietic tree, indicating the diversity of AML. The patients with leukemia cells projecting into the same hematopoietic lineage showed similar features, while patients with leukemia cells projecting into the same different hematopoietic lineage showed much different features. Therefore, Projection of leukemia cells into hematopoietic tree is an accurate and unbiased approach for leukemia classification. Furthermore, we found the patient with leukemia cells projecting to the root of the hematopoietic tree had the worst results.

### Hypothesis about the origin of leukemia heterogeneity

Pseudotime inference of leukemia cells further showed that the trajectories of leukemia cell were similar to that of their healthy counterparts (**Extended Data Fig. 5f**). In this way, we could assume that leukemia cell subpopulations are not a bunch of independent subpopulations but are a series of lineage-related cells in leukemogenesis. Therefore, we hypothesize the mutant leukemia initial cells or progenitors partially maintain its original developmental trajectory and terminate development at different stages due to loss of different functional genes, leading to a serial of dysfunctional leukemia cell subpopulations, instead of developing into a homozygous population.

### Cell population dynamics of BMMCs in patient ALL3 and the underlying genes

Patient ALL3, a 4 year old boy, was diagnosed with B-precursor ALL in Blood Diseases Hospital, Chinese Academy of Medical Sciences (CAMS)/ Peking Union Medical College (PUMC) in Nov 2012 ^35^. The patient was refractory to a prolonged chemotherapy, and later achieved remission after Imatinib treatment. The patient relapsed and no longer responded to Imatinib treatment 5 months after remission (**Fig. 5a**). scRNA-seq data of BMMCs at diagnosis (ALL3.1), refractory (ALL3.2), remission (ALL3.3) and relapse (ALL3.4) were generated, which allowed us to investigate leukemia progression and the underlying mechanisms. By integrating healthy BMMCs and HSPCs as reference, leukemia cells could be easily distinguished from normal cells since leukemia cells are patient specific and normal cells are shared by all samples. Cells from healthy reference formed a notched circle on the tSNE projection (**Fig. 5b, 5c**), while leukemia cells formed several distinct clusters at the breach or inside of the notched circle (**Fig. 5b**). At diagnosis, leukemia cells account for 87.4% of total BMMCs and form a major cell cluster surrounding with some minor cell subpopulations (**Fig. 5d**). After prolonged chemotherapy, the percentage of leukemia cells in BMMCs slight decrease while the size of some surrounding minor cell subpopulation relatively increased (**Fig. 5e**). After Imatinib treatment, leukemia cells almost completely disappeared (∼1.9%) during remission (**Fig. 5f**). However, the leukemia cells come back and become dominant (∼82.7%) after relapse, with a significant reduction of normal cells (**Fig. 5g**). Especially, leukemia cells before and after relapse were projected to different coordinates of tSNE projection (**Fig. 5b, 5g**), indicating the relapsed leukemia cells are quite different from the leukemia cells in the early stage. The relapsed leukemia cell has the highest fraction of cells in active cell cycle and has the highest entropy among all cell clusters (**Fig. 5h, 5i and Extended Data Fig. 6a**), indicating the proliferation and transcriptional complexity of leukemia cells significantly increased after relapse. In summary, these results vividly showed the pronounced dynamics of leukemia cells during clinical treatments and relapse.

**Fig. 5.**
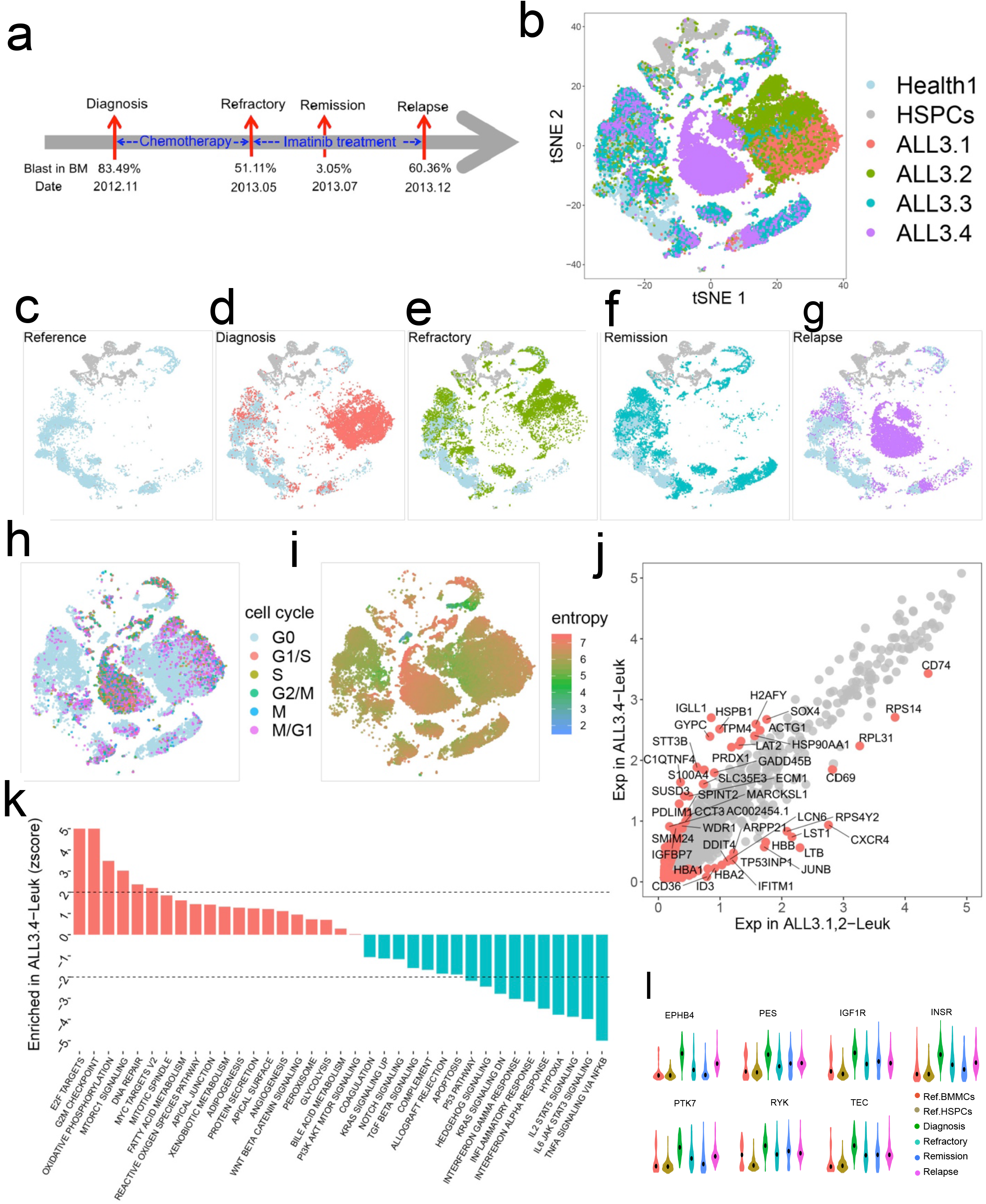
Cell population dynamics of cell populations in patient ALL3 and the underlying genes. a. Sampling information the patient ALL3 at diagnosis, refractory, remission and relapse. b. tSNE projection of all BMMCs from this patient and reference cells. c-g. tSNE projection of reference cells and BMMCs at each phase. Reference cells only (c), diagnosis (d), refractory (e), remission (f) and relapse (g). h. Cell cycle of patient BMMCs and reference cells. i. Entropy of patient BMMCs and reference cells. j. Significantly differential genes between pre-and post-relapse leukemia cells. k. Significantly enriched pathway and gene sets between pre- and post-relapse leukemia cells. l. The expression dynamics of some tyrosine kinase (TKs) based on scRNA-seq.

According to the above observation, we could conclude that leukemia cells experienced dramatic changes across time. Identifying the differentially expressed genes and pathways between pre- and post-relapse leukemia cells could enhance our understanding of relapse process. We identified 243 significantly upregulated genes and 79 significantly downregulated genes (fold change >2) after relapse (**Fig. 5j**). The upregulated genes include H2AFY, IGLL1, GYPC, HSPB1, STT3B, C1QTN4, SUSD3 and PDLIM1 (**Fig. 5j**), which significant enriched in E2F targeted genes, G2M checkpoints, oxidative phosphorylation, MTORC1 signaling and so on (**Fig. 5k**). These gene sets have been reported to promote the cell proliferation in tumor, which is consistent with our observation that relapsed leukemia cells expressed the highest level of cell cycle genes and have the highest entropy. The 79 downregulated genes include CXCR4, DUSP1, JUNB, LST1, LTB, RPS14, RPL31 and RPS4Y2 (**Fig. 5j**), which significant enriched in IL-6 JAK STAT3 signaling, TNFA signaling pathway, IL-2 STAT5 signaling, interferon alpha response, hypoxia, inflammatory response, interferon gamma response, KRAS signaling pathway and hedgehog signaling and so on (**Fig. 5k**). Imatinib inhibits the enzymes activity of tyrosine kinase that has been shown to play a central role in the pathogenesis of human cancers. We observed the expressions of a lot of tyrosine kinases were changing during the treatment and relapse (**Fig. 5l**). In summary, relapsed leukemia cells showed substructure shift and molecular difference with leukemia cells before relapse.

### Leukemia progress model for patient ALL3

After analyzing the dynamics of BMMCs, we zoomed in leukemia cell subpopulations to provide biological insight on leukemia progress. The total leukemia cells from the 4 time points were classified into 6 subpopulations (**Fig. 6a**). Using CCI, we found the counterparts of the 6 leukemia cell subpopulations were B cell progenitors, namely pro-B I, pro-B II, pre-B and immature-B (**Fig. 6b**). The major leukemia subpopulations at diagnosis and refractory (clusters C1 and C2) were pre-B-L and immature-B-L, while the major relapsed leukemia cells (clusters C5 and C6) were pro-B-L (**Fig. 6b**). Since pro-B is the progenitor of pre-B and pre-B is the progenitor of immature B in heathy hematopoietic lineage, we could assume that relapsed leukemia cells have increased stemness and stronger differentiation potential than leukemia cells at early stage, which is consistent with our observation that relapsed leukemia cells were in high proliferation states with the highest cell cycle activity and the highest entropy (**Fig. 5h, 5i**).

**Fig. 6.**
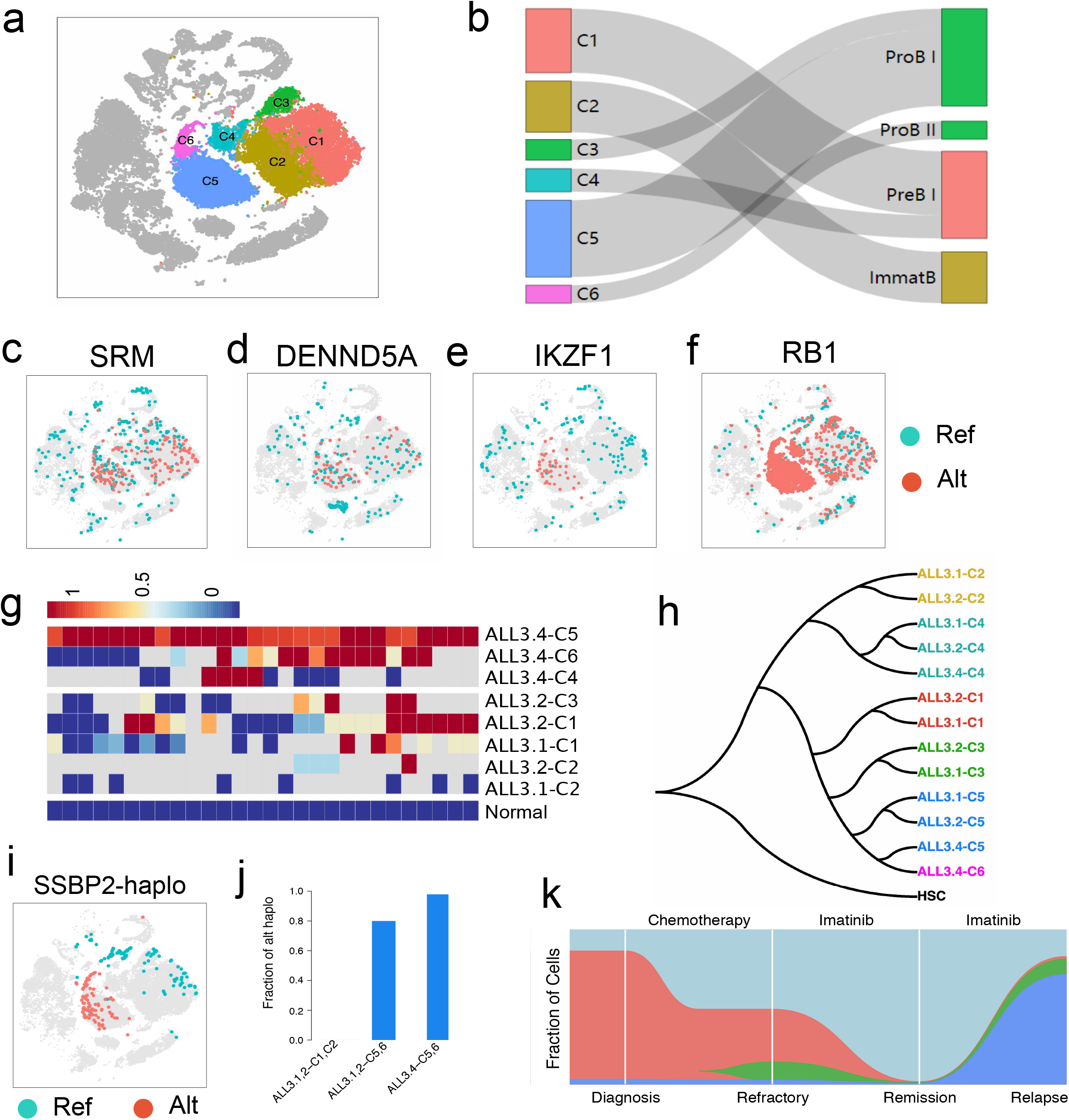
Tracing relapsed leukemia cells and leukemia progression model. a. Clustering of leukemia cell subpopulations of ALL3. b. Healthy counterpart of leukemia cell subpopulation, the healthy cell populations are listed by order of lymphoid lineage. c-d. Distributions of leukemia cell specific high variants on SRM (C) and DENND5A (D). e-f. Distribution of relapsed leukemia cell specific high variants on IKZF1 (e) and RB1 (f). g. Distribution of genetic variants in different cell subpopulations. h. Hierarchical tree of leukemia cell subpopulations, in which relapsed leukemia cells and earlier leukemia cells in C5 were clustering together. i. Distribution of reference haplotype and alterative haplotype of SSBP2 in BMMCs. j. Frequencies of relapsed-specific SSBP2 haplotype (alt haplo) in different leukemia cell subpopulations. k. Progression model of patient ALL3.

We identified single cell transcriptomic variants at different phases and found increased variants in relapsed leukemia cells (**Extended Data Fig. 6b**), consistent with the expectation that mutations are progressively accumulated during cancer progress ^36^. We identified the leukemia cell or its subpopulation specific variants by comparing the variants between different cell populations. Among the variants that are significantly different between leukemia cells and normal cells, chr1:11055657;T>C on SRM and chr11:9173222;A>C on DENND5A are two mutations distributed in all leukemia cell subpopulations and might play an important role in in early leukemogenesis (**Fig. 6c, 6d**). On the other hand, the mutational alleles on IKZF1 (chr7:50382590, G>A, p.G158S) and RB1 (chr13:48409776, T>C) are extremely concentrated in relapsed leukemia cells (**Fig. 6e, 6f**), indicating these mutations raised during late phase of leukemia progress. Especially, IKZF1^G158S^ is a deleterious mutation associated with leukemia and is a predictor of poor outcome in ALL ^37,38^, potentially indicating that IKZF1^G158S^ mutation might play an functional role in the relapse of the patient.

We further separate each leukemia cell cluster into multiple subpopulations according to time points of data collection. We found the transcriptomic mutations was accumulating during the leukemia progress (**Fig. 6g**). We further constructed a hierarchical tree based on correlation coefficient of gene expression between pairwise subpopulations, choosing HSCs as root of the tree. The results showed cell subpopulations from different time points in the same cluster located on the same branch (**Fig. 6h**). The major relapsed leukemia cells (ALL3.4-C5, ALL3.4-C6) and a minor leukemia cell subpopulation from early phase (ALL3.1-C5 and ALL3.2-C5) are located on the same branch (**Fig. 6h**), implicating the two subpopulations are similar to each other and potentially shared a common progenitor. In this way, we hypothesize that the relapsed leukemia cells were derived from this minor leukemia subpopulation in early phase (ALL3.1-C5 & ALL3.2-C5) that developed resistance to Imatinib and rapidly expanded to the major subpopulations in relapse. SSBP2, a tumor suppressor gene involved in the maintenance of genome stability ^39^, contains two haplotypes in the leukemia cells, namely reference haplotype and alternative haplotype. Interestingly, reference haplotypes concentrated in early leukemia cells and normal cells, while alterative haplotypes exclusively enriched in cluster the major relapse leukemia cells (C5 and C6) (**Fig. 6i**). We found the fraction of cells with alternative haplotype in early leukemia cells (ALL3.1-C5 and ALL3.2-C5) was nearly as high as that in relapsed leukemia cells (ALL3.4-C5) (**Fig. 6j**), which strongly supports the notion that relapsed leukemia cells originated from the early minor leukemia cell subpopulation. Furthermore, the genetic variants in leukemia cells subpopulations support that relapsed leukemia cells were derived from minor leukemia cell populations from early phase (**Extended Data Fig. 6d, 6e**). Based on these observations, we proposed a model for leukemia progress (**Fig. 6k**), which vividly showed the dynamics of leukemia cells during diagnosis, refractory, remission and relapse.

## Discussion

Distinct models have been proposed to explain the origin of leukemia heterogeneity ^32–34^, However, these studies usually only focus on analyzing relationships among leukemia cell subpopulation thus lost the whole picture of leukemogenesis. It has the potential to provide novel insight by integrating both normal cell and leukemia cell subpopulation for exploring their potential relationships. However, it is big challenge to identify the healthy counterparts of each leukemia cell subpopulation based on a single signature due to complex leukemogenic process. Consistent supports from multiple signatures could generate more reliable results because different measurements provide complementary information about the relationship between cell subpopulation. We developed CCI that integrates multiple measurements for searching the healthy counterpart of each leukemia cell subpopulation in reference hematopoietic cells. Interestingly, the healthy counterparts of the leukemia cell subpopulations within each patient were closely adjacent in hematopoietic lineages, indicating leukemia cell subpopulations with lineage-relationship.

Furthermore, identification of the healthy counterparts of leukemia cells not only facilitate our understanding of the leukemia progression and leukemia heterogeneity, but also could provide biological insights on the features and functions of the leukemia cell subpopulation via their well-annotated healthy counterparts. Finally, identification of the healthy counterpart of leukemia cell subpopulations have a lot clinical implication such as prediction of leukemia subtype and clinical outcome.

Analyses of clinical data with multiple time points have the potential to provide details about leukemia progressions. Our analyses of patient ALL3 with data at 4 time points, namely diagnosis, refractory, remission and relapse, vividly showed dynamics of cell population shifting during treatment and relapse. Variants calling from single cell transcriptomes identified the leukemia specific mutations and relapsed leukemia cells specific mutations that are potentially associated with leukemia progressions. Both gene expression and mutations of leukemia cells support that relapsed leukemia cells originated from an early minor leukemia cell population. We further proposed a leukemia progress model, in which the leukemia cells at the diagnosis are similar to malfunction immature B or pre-B. Majority of leukemia cells was killed after Imatinib treatment and patient achieved remission, However, a minor leukemia cell subpopulations, with the highest similarity to malfunction pro-B, developed Imatinib resistance and later rapid expansion lead to relapse. In summary, our study vividly showed that leukemia heterogeneities are associated with cancer progression and therapy outcomes.

## Methods

### Clinical samples

In total, 18 BMMCs samples and 1 PBMC sample were analyzed in this study, with detail information (**Table S1**). Among them, 14 samples were collected in Blood Diseases Hospital, CAMS/PUMC and Nanfang Hospital, Southern Medical University. All individuals signed an informed consent form approved by the IRB of the Blood Diseases Hospital, CAMS/PUMC and Southern University of Science and Technology (SUSTech). The other 4 BMMCs samples with scRNA-seq data were collected from Zheng et al. ^15^. The diagnosis of leukemia was established according to the criteria of the World Health Organization ^40^. Overall, there are 5 healthy samples, 7 AML samples, 6 ALL samples from 3 patients, among which the patient ALL3 were sampled at diagnosis, refractory, remission and relapse.

### Cell preparation and flow cytometry

BMMCs were isolated from whole bone marrow aspirate by Ficoll density gradient separation (GE Healthcare), resuspended in 90% FBS + 10% DMSO, and cryopreserved in liquid nitrogen. To prepare cells for FACS, frozen BMMCs vials were thawed in a 37°C water bath for 2 mins. Vials were removed once only a tiny ice crystal was left. Thawed BMMCs were washed and resuspended in 1 PBS and 20% FBS. After recovery, FACS sorting was performed on a Becton Dickinson FACSAria II (BD Biosciences, Denmark) to remove the dead cells. BMMCs were stained with pre-conjugated CD34-PE antibody for 15 min on ice. Non-specific binding was blocked by incubation in FACS buffer (Life Technologies). The unbound antibodies were removed using 5 ml wash buffer. The CD34+ cells were sorted out by FACS Aria™ II. The final concentration of thawed cells was 1X 10^6^ cells per ml.

### Single cell library preparation and sequencing

The single cell RNA sequencing libraries of BMMCs and CD34+ cells from healthy individuals and leukemia patients were generated using 10X genomics. In brief, cell suspensions were loaded on a 10x Genomics Chromium Single Cell Instrument (10x Genomics, Pleasanton, CA) to generate single cell GEMs. Approximately 20,000 cells were loaded per channel. Single cell RNA-seq libraries were prepared using the Chromium Single Cell 3^’^ Gel Bead and Library Kit (P/N 120237, 120236, 120262, 10x Genomics) following the protocols ^15^. Sequencing libraries were loaded at 2.4 pM on an Illumina HiSeq4000 or Illumina NovaSeq 6000 with 2 × 75 paired-end kits.

### Alignment, demultiplexing, UMI Counting and normalization

The Single Cell Software Cell Ranger Suite 2 was used to perform reads alignment, barcode demultiplexing, transcripts assemblies and expression counting (https://support.10xgenomics.com). The gene-cell barcode matrix was filtered based on number of genes detected per cell (any cells with less than 500 or more than 4000 genes per cell were filtered out), and percentage of mitochondrial UMI counts (any cells with more than 10% of mitochondrial UMI counts were filtered out). UMI normalization was performed by first dividing UMI counts by the total UMI counts in each cell, followed by multiplication with the median of the total UMI counts across cells. Then, we took the natural log of the UMI counts.

### Dimension reduction and clustering analysis

Dimension reduction was performed by PCA and tSNE ^41^, as described in Macosko et al^4^. The highly variable genes were inferred based on normalized dispersion following Macosko et al. ^4^. The top 30 principal components (PCs) were chosen for tSNE and clustering analysis, according to the explained variances. Single cell clustering was performed by k-nearest neighbor (KNN) graph and Louvain algorithm. Raw clusters with few differential expressed genes were merged to avoid excessive classification.

### Inferences of hematopoietic lineages and lineage coordinated genes

Hematopoietic lineages and cell pseudotime of each hematopoietic lineage was inferred by Slingshot ^19^, in which PCA was implemented and cell clusters were predefined. The cell clusters inferred by KNN graph were used as input for Slingshot to infer cell trajectories. The constructed hematopoietic lineages was visualized by a force-directed layout in SPRING ^20^, in which dimension reduction were generated by DiffusionMap ^42^. Differentially expressed genes along the pseudotime were identified using negative binomial tests in Monocle2 ^7,43^, with the smoothing parameter set to three degrees of freedom. For generating heatmaps of gene expression dynamics, the normalized UMI counts were log transformed and smoothed using Loess regression with the degree of smoothing (span) set to 0.75. Heatmaps for gene expression profile clustering were generated using heatmap function in R. While the other graphics were generated using ggplot2 in R.

### Gene regulatory network (GRN) score

The GRN score ^23^ reflect the regulator-target relationships in the context of trajectory progression, which ranks transcriptional regulators based on their correlation with the trajectory, the correlation with their predicted targets, and the extent to which target genes are regulated during the trajectory, which defined as below,

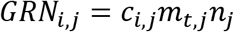

where *GRN_i,j_* is the GRN score for regulator *i* along trajectory *j*. *c_i,j_* is the mutual information between transcriptional regulator *i* and trajectory *j*, *m_t,j_* is the average mutual information between predicted target gene *t* and trajectory *j*, *n_j_* is the number of predicted targets regulated along trajectory *j*.

### Lineage coordinated TF networks

Human transcription factors (TFs) were downloaded from TcoF-DB v2 ^44^. Lineage coordinated TF networks were constructed following Fletcher et al. ^45^. In short, TFs showing significantly gene expression changing along inferred lineage are lineage coordinated TFs. The co-expression networks of lineage coordinated TFs was called lineage coordinated TF networks. A lineage coordinated TF was added into the lineage coordinated TF networks if it correlated with at least 5 other lineage coordinated TFs (Pearson correlation coefficient > 0.1). TFs correlation heatmaps were generated with NMF R package.

### Gene sets enrichment analysis (GSEA)

GSEA determines whether a priori defined set of genes shows statistically significant differences between two biological states. The original GSEA was developed for gene-expression assays of bulk data ^27,46^, which may lost accuracy when directly implement on scRNA-RNA data. In order to take advantage of thousands of single cell transcriptomes, we designed an approach as below: (1) Gene expression was averaged from 30 random cells from each cluster due to the high dropout rates of scRNA-seq data. Genes were ranked according to their expression level for each set of cells. (2) Recovery curve was created by walking down the gene list, and steps were increased when we encounter a gene in the gene set. Area under the curve (AUC) was computed as the indicator of enrichment for a certain gene set. And only AUC of top 5000 ranked genes was considered. (3) To compare the different enrichment of two cell populations for a gene set, we calculate the Zscore of AUC in one cell population relative to the AUC distribution in another cell population. Gene sets from Msigdb3.0 (Molecular Signatures Database) were used for analysis by our approach.

### Counterpart composite index (CCI)

It is interesting to systematically compare the developmental trajectories of leukemia cells with that of healthy hematopoietic cells. However, searching the counterpart of a leukemia cell subpopulation in healthy hematopoietic cells is very challenging due to the significant difference between leukemia cells and healthy hematopoietic cells. Based on simulated data, our initial analyses showed that different approaches may produce different results. However, different statistics have much higher probability to generate consistent result when two cell populations have counterpart relationship than these when two cell populations have uncertain relationship. Thus, the composite likelihood of the statistics is the highest when two cell populations have counterpart relationship. Here, we developed an approach, called counterpart composite index (CCI), to search for healthy counterpart of each leukemia cell subpopulation by integrating multiple statistics to project leukemia cells into healthy hematopoietic cells. CCI integrates Euclidean distance of gene expression, correlation of gene expression, weighted distance in top PCs and difference of gene set enrichment for composite likelihood statistical framework as below:

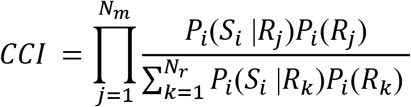

In which, *S_i_* is the score of measurement *i*, *R_j_* is the candidate reference cell population *j*, *N_r_* is the number of reference cell population, *N_m_* is the number of composite measurements in CCI.

The computational details of each statistics were described as follows:

1. Euclidean distance of gene expression Considering the prevalent of dropouts and sample size asymmetry, gene expression level was averaged from 30 random cells from each population. The sampling was repeat by ten thousand times for each pairwise populations, from which the median value of expression difference was used as the statistical measurement of two populations. In the meanwhile, pairwise statistical distribution was built from the sampling and computations.
2. Correlation of gene expression Spearman’s rank correlation coefficient between cell populations was calculated to measure the similarity of gene expression. The sampling and distribution construction were following the same strategy as above.
3. Weighted distance in top PCs The top principal components, which could represent the distance between cell populations or even single cells, were integrated into CCI. Euclidean distance in embedding space defined by top 30 eigenvectors was weighted by their explained variances.
4. Difference of gene set enrichment It is well known that cells with similar function have similar biological process and show similar gene set enrichment. Recovery curve of each gene set was constructed and AUC value of top 5000 ranks were used for the indication of cell similarity. Averaged gene expression from 30 random cells of each population were used to calculate the responding AUCs of 50 hallmark gene sets from Msigdb. Median value and statistical distribution were derived from ten thousand of samplings.

Facilitated by the comprehensive cell atlas of BMMCs and HSPCs, we could identify the healthy counterparts of each leukemia cell subpopulation in HSPCs using CCI, which could greatly benefit our understanding of the progression of leukemia.

### Identifying the genetic variants in single cell transcriptome

There are a few reads in each locus of the single transcriptomic data; thus, it is very difficult to directly detect the cell specific variants. In order to reliably detect single-cell expressed specific variants, reads from every single individual were pooled to do variants calling. An in-house script was used to assign each single cell with its associated variants, by checking the variant on indexed reads. To reduce the artifacts and false positives, we used following criteria to filter the cell variants: 1) not missing in at least 20 cells; 2) at least 5 cells with 2 reads sequenced; 3) variants observed in at least 3 cells. We found that the number of detected transcript mutations in each leukemia patient was much higher than that in each healthy individual, partially due to increased number of expressed genes in leukemia cells. The single nucleotide variations (SNVs) identification could be affected by the copy number variations (CNVs) change in studied regions. We identified genome wide CNVs in leukemia cells, which could affect the genome-wide identification of SNVs. Therefore, we used the deletion regions from the bulk data to filter the covered homozygous SNVs, as well as the associated signals. CNVnator (v0.3.2) was used to call CNVs from the whole-genome sequencing data, with a depth of ∼30X, of ALL3. The somatic CNVs was then generated by comparing CNVs at different time points to CNVs detected from its saliva sample.

### Haplotype tracing

We observed a sequential of homozygous variants of SSBP2 from the major relapsed leukemia cell subpopulation ALL3.4-C5, which is the alternative haplotype of SSBP2. We analyzed 39 polymorphisms in SSBP2 to investigate the distribution of the alternative haplotype in different cell populations. Any cell with more than 5 loci showing the same alleles as that in alterative haplotype (P<0.05) were thought contained the alterative haplotype.

## Acknowledgements

This study was supported by National Key R&D Program of China (2018YFC1004500), National Science Foundation of China (81872330, 31741077), The Shenzhen Science and Technology Program (JCYJ20170817111841427, KQTD20180411143432337). We thank Xi Chen for discussion and editing of the manuscript. The funders had no role in study design, data collection and analysis, decision to publish, or preparation of the manuscript.

## Author contributions

W.J. conceived this project. Y.P., Y.Z., Q.L. and X.Z. collected the samples. Y.P. did the experiments. P.Q., W.H., R.F., X.W. and G.M. developed computational approach and performed the data analysis. W.J., C.T. and N.H. supervised this project. W.J. Q.L. and N.H. wrote the manuscript with inputs from all authors.

## Competing interests

The authors have declared that no competing interests exist.

## Availability of sequencing data

The raw sequence data reported in this paper have been deposited in the Genome Sequence Archive in BIG Data Center, under accession numbers HRA000084.

## Supplementary Data are available online

**Tables S1. The samples information.**

**Extended Data Fig. 1. Basic information of scRNA-seq data and the cell subpopulations in BMMCs.**

a-c. Basic data information. Number of cells (a), number of genes (b) and number of UMI (c) after data quality control in all samples.

**d.** Differentially expressed genes in two distinct plasma cell populations.

**e.** The major Cell types in 4 healthy BMMCs.

**f.** T cell subpopulations and NK cells in healthy BMMCs.

**g-h**. Comparison of cell types in healthy BMMCs (g) and PBMCs (h).

**i-j**. Comparison of cell activity and proliferation in healthy BMMCs and PBMCS revealed by entropy (i) and cell cycle (j).

**Extended Data Fig. 2. Single cell profiles of healthy BMMCs and HSPCs.**

**a.** Isolation of CD34+ cells by FACS sorting.

**b.** Normalized expression level and expression percentage of cell type specific genes in HSPCs subpopulations.

**c-d**. Cell activity and proliferation in healthy BMMCs revealed by cell cycle (c) and entropy (d) analysis.

**e-f**. Cell activity and proliferation in healthy HSPCs revealed by cell cycle (e) and entropy (f) analysis.

**g-h**. Comparison of cell activity and proliferation in healthy BMMCs and HSPCs revealed by cell cycle (g) and entropy (h) analysis.

**Extended Data Fig. 3. Cell atlas and hematopoietic lineages of cells from HSPCs and BMMCs.**

**a.** Trajectory inference of HSPCs by slingshot.

**b.** tSNE projection of cells from HSPCs and BMMC, colored by HSPCs and BMMCs, respectively.

**c.** tSNE projection of cells from HSPCs and BMMC, colored by inferred cell types.

**d.** Hematopoietic lineages were re-inferred by incorporating some BMMC cell populations, which were essentially consistent with that inferred by HSPCs.

**e.** The HSPCs were located on the core of hematopoietic lineages, while BMMCs (red) extended hematopoietic lineages at branch end.

**Extended Data Fig. 4. Lineage coordinated genes, TFs and TF networks underlying DC lineage, neutrophil lineage, monocyte lineage, Ba/Eo/Ma lineage, Mk lineage and Ery lineage.**

a-b. Heatmap of transcriptomic dynamics in DC lineage (a) and Ery lineage (b).

c-h. The top 10 coordinated TFs, correlation heatmap of coordinated TF, connectivity graph of coordinated TFs, dynamics of TFs expression in regulatory networks at different stages with DC lineage (c), neutrophil lineage (d), monocyte lineage (e), Ba/Eo/Ma lineage (f), Mk lineage (g) and Ery lineage (h).

**Extended Data Fig. 5. Features and pseudotime of leukemia cells.**

**a.** Shared high expressed genes among the leukemia patients.

**b.** Boxplot of expressed cell cycle genes in each cell in healthy BMMCs and leukemia BMMCs.

**c-d**. Cell cycle pattern of leukemia BMMCs (b) and healthy BMMCs (c).

**e.** G2M/G1S ratio in healthy and leukemia BMMCs.

**f.** Pseudotime of leukemia cells.

**Extended Data Fig. 6. Dynamics of leukemia cells and mutation profile of ALL3 at diagnosis, refractory, remission and relapse.**

**a.** Distribution of expressed cell cycle genes at 4 stages of ALL3.

**b.** Potential mutation carriage of each cell at 4 stages of ALL3.

**c.** Venn diagram of detected mutations at the 4 time points.

**d.** Alleles and haplotypes of SSBP2 at different cell subpopulations in ALL3.

**e.** Hierarchical relationship of leukemia cell subpopulation revealed by transcriptional mutations.

