## Extended Fig1-6 and TableS1 for "Integrated decoding hematopoiesis and leukemogenesis at single-cell resolution and its clinical implication"

Extended Data Fig. 1

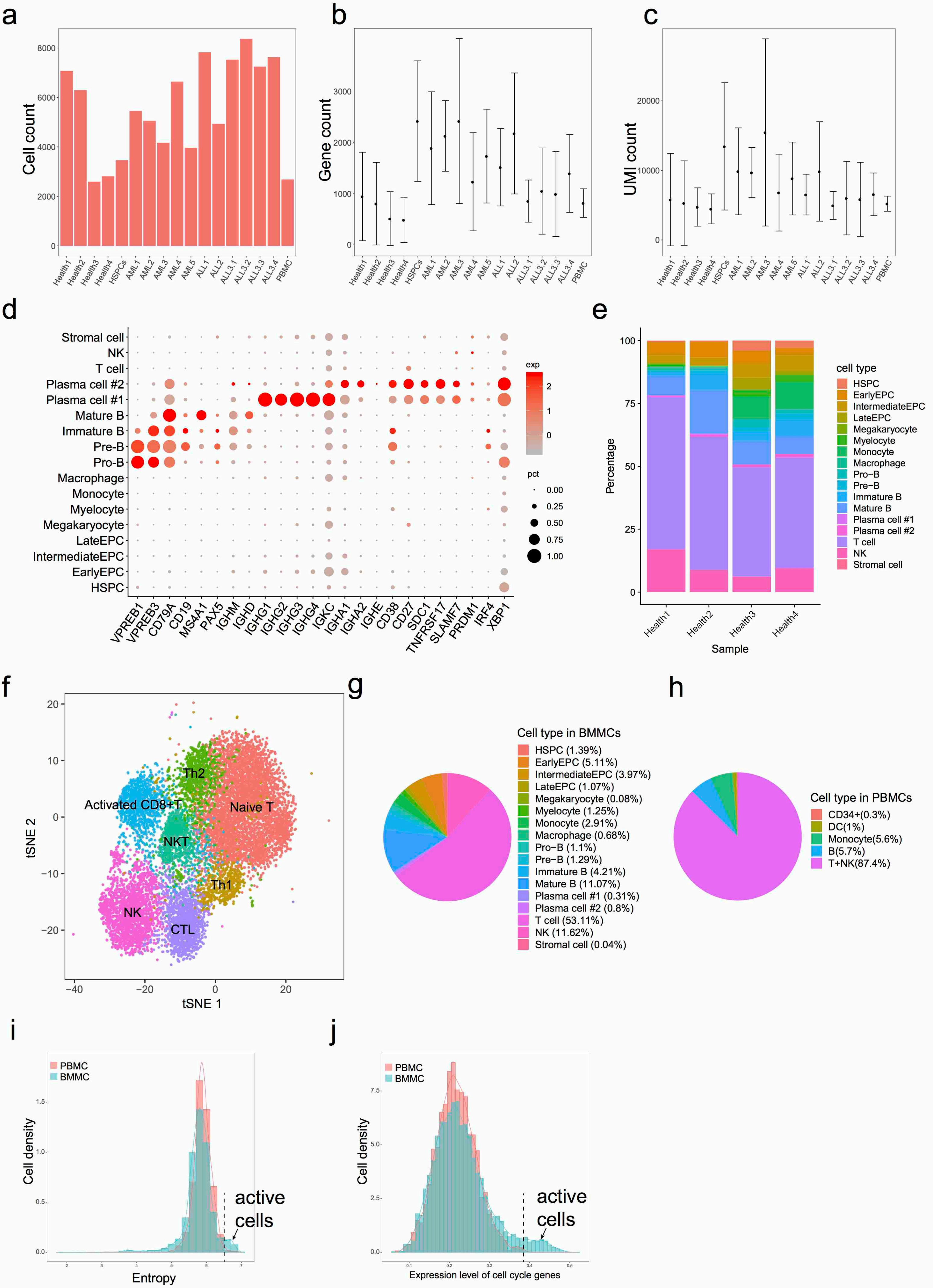

Extended Data Fig. 2

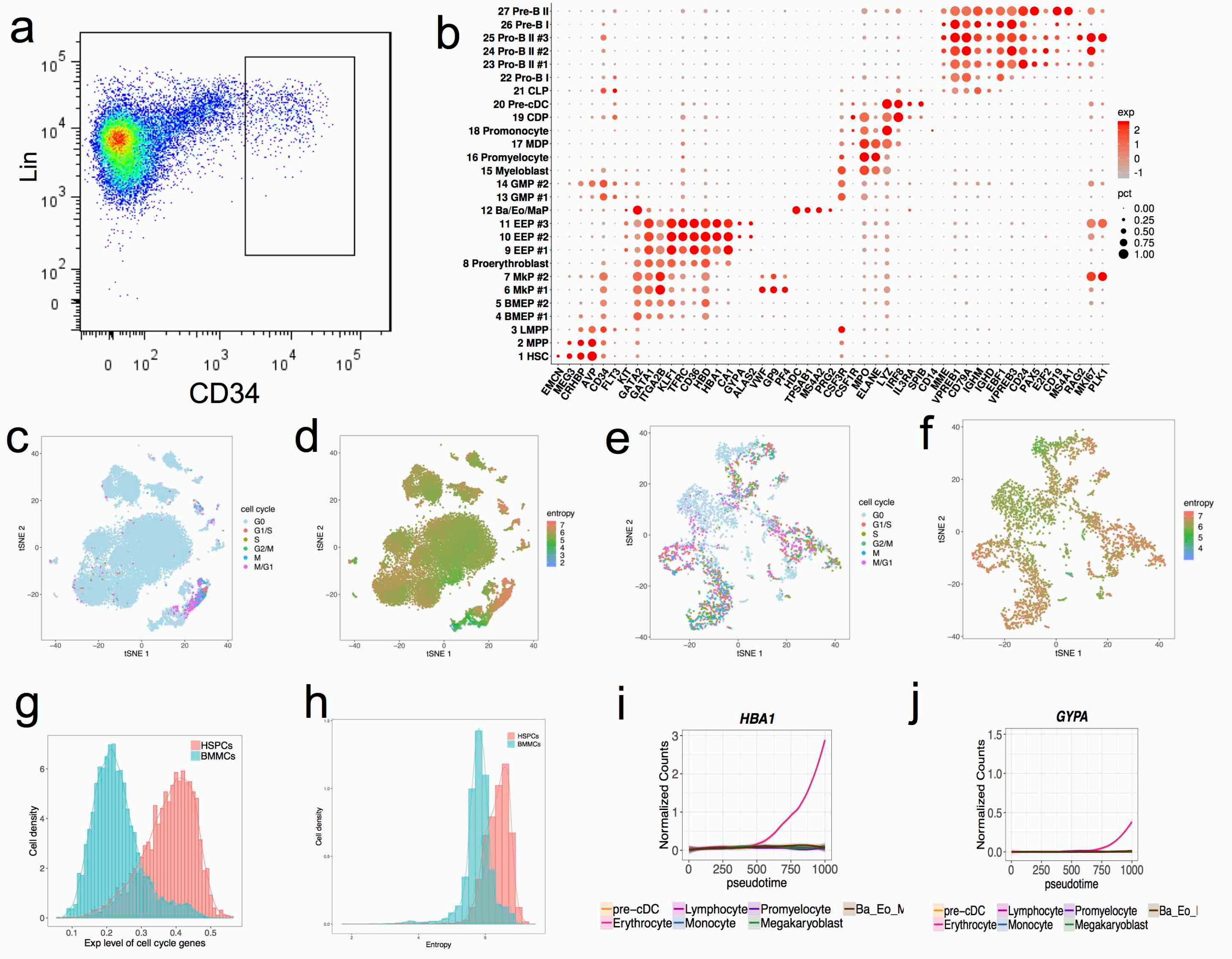

Extended Data Fig. 3

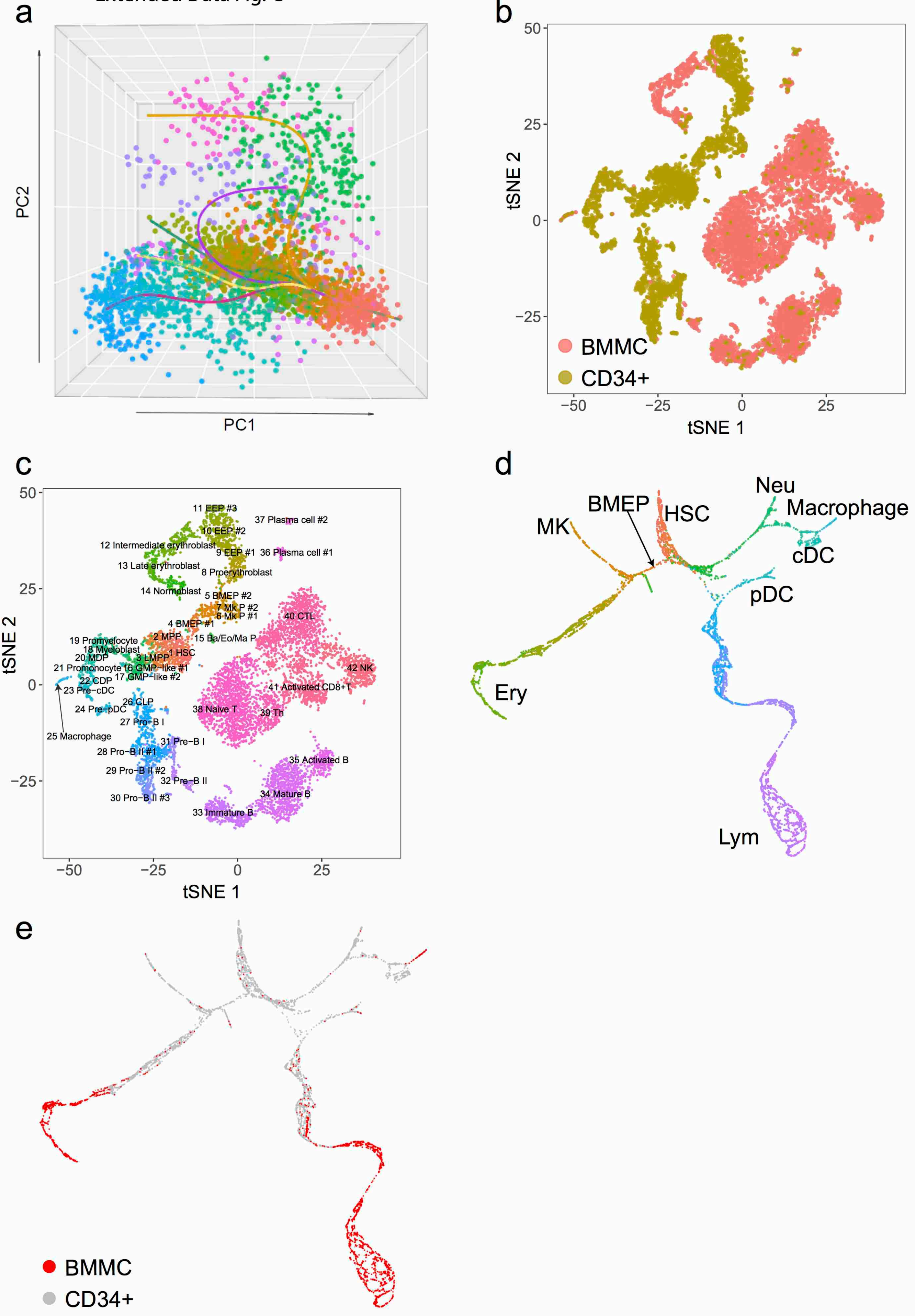

Extended Data Fig. 4

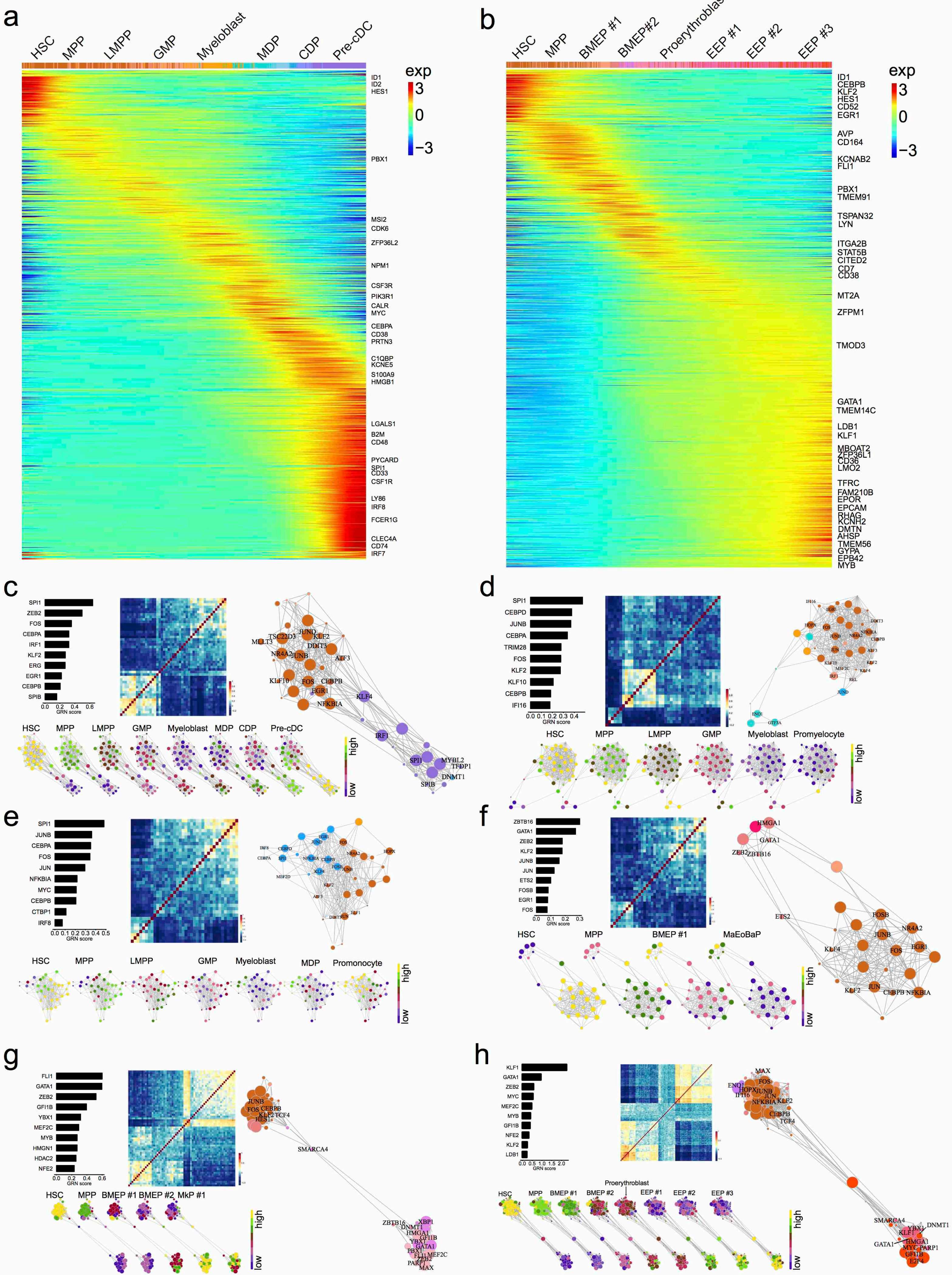

Extended Data Fig. 5

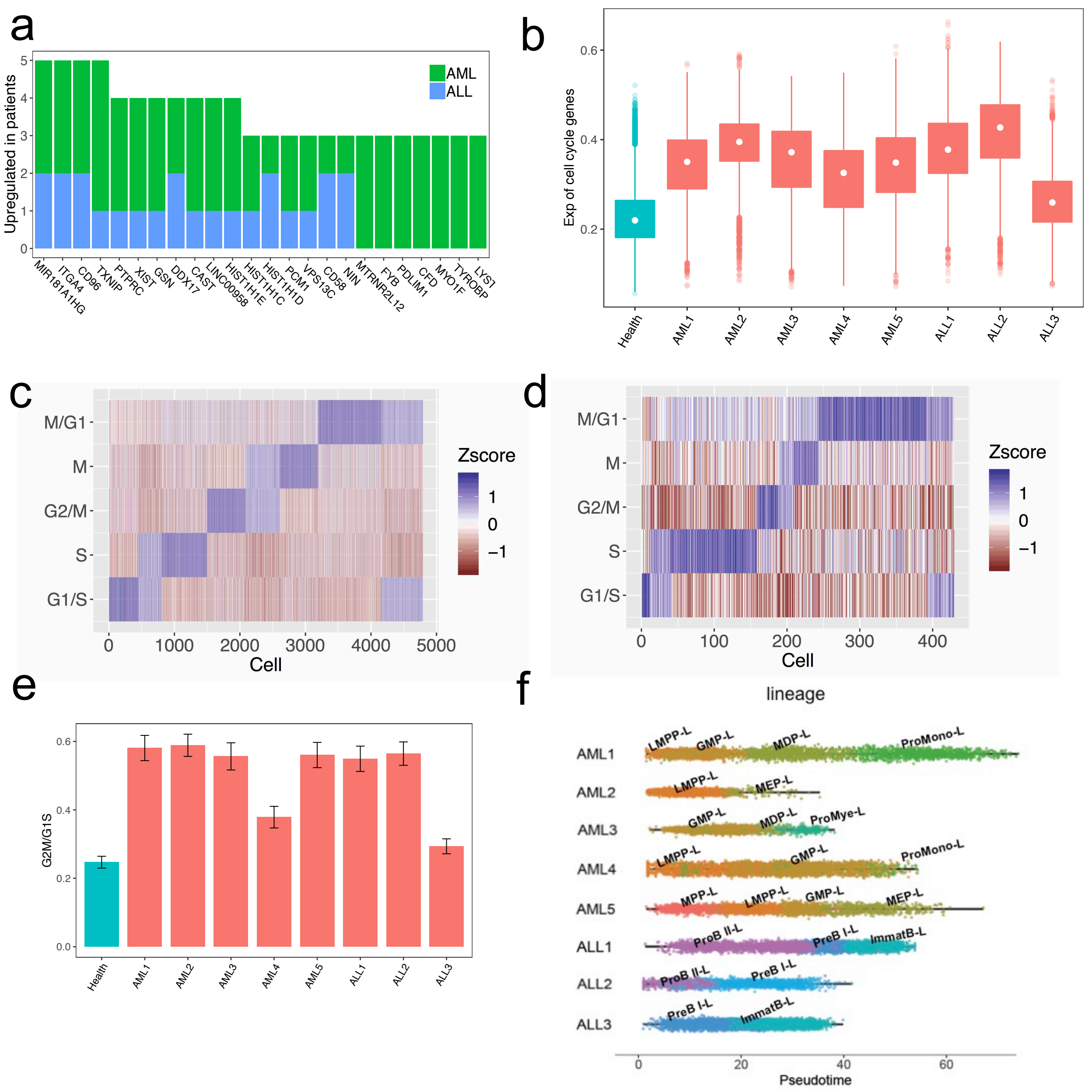

Extended Data Fig. 6

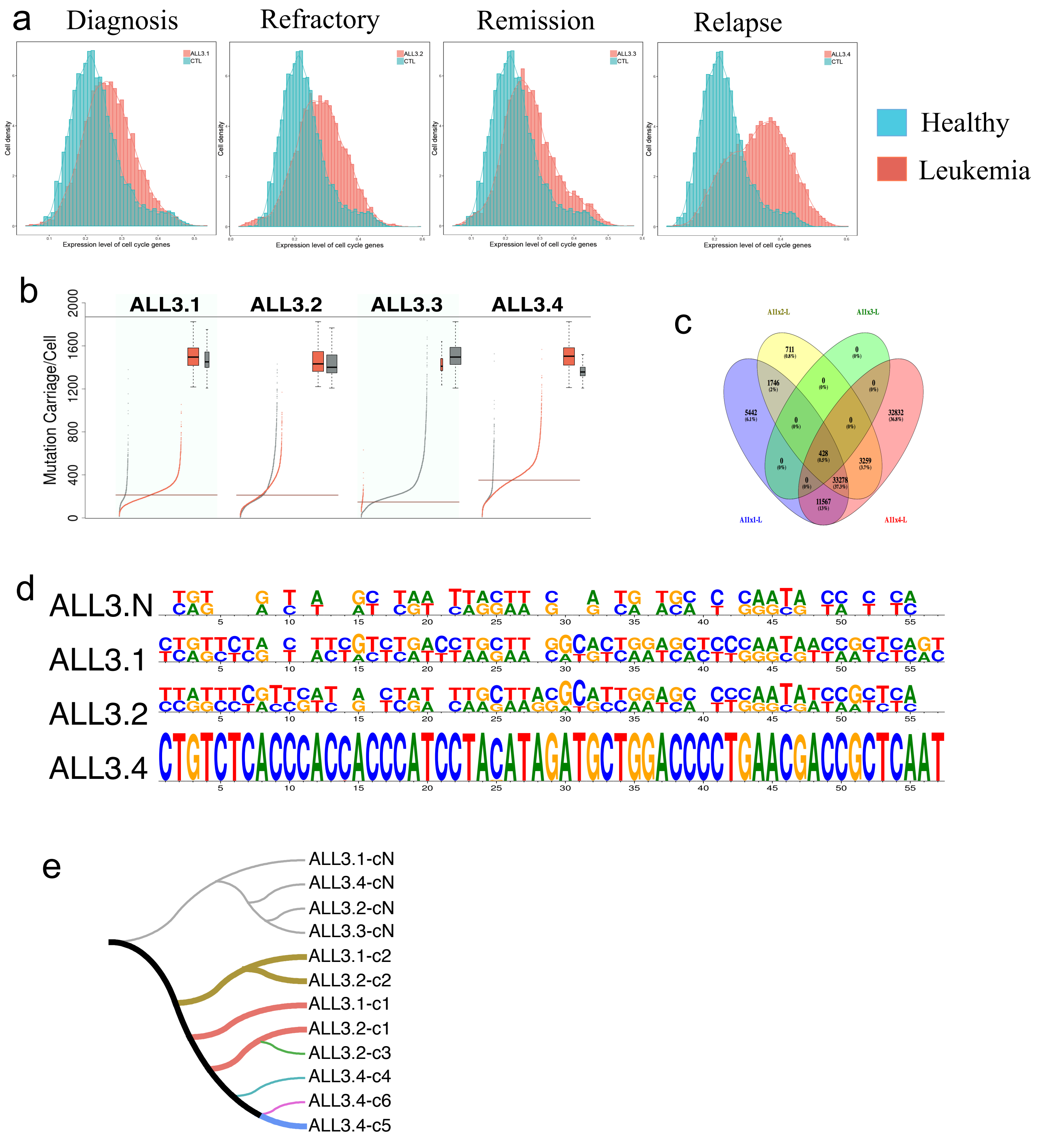

**Table S1**. Sample information

| Sample ID | Clinical diagnosis | Sampling time | Cell type | Cell Number |
|---|---|---|---|---|
| AML1 | AML-M4 | Diagnosis | CD34 | 5,457 |
| AML2 | AML-M2 | Diagnosis | CD34 | 5,059 |
| AML3 | AML-M2 | Diagnosis | CD34 | 4,168 |
| AML4 | AML-? | Diagnosis | BMMC | 6,638 |
| AML5 | AML-M5 | Diagnosis | CD34 | 3,967 |
| ALL1 | ALL-B | Diagnosis | BMMC | 7,827 |
| ALL2 | ALL-B | Diagnosis | CD34 | 4,933 |
| ALL3.1 | ALL-B | Diagnosis | BMMC | 7,526 |
| ALL3.2 | ALL-B | Refractory | BMMC | 8,370 |
| ALL3.3 | ALL-B | Remission | BMMC | 7,247 |
| ALL3.4 | ALL-B | Relapse | BMMC | 7,628 |
| AML027 | AML | Diagnosis | BMMC | 2,977 |
| AML035 | AML | Diagnosis | BMMC | 4,828 |
| HSPCs | Healthy | Donor | CD34 | 3,465 |
| Health1 | Healthy | Donor | BMMC | 7,074 |
| Health2 | Healthy | Donor | BMMC | 6,299 |
| Health3 | Healthy | Donor | BMMC | 2,593 |
| Health4 | Healthy | Donor | BMMC | 2,814 |
| PBMC | Healthy | Donor | PBMC | 2,685 |
